# Regulatory elements can be essential for maintaining broad chromatin organization and cell viability

**DOI:** 10.1101/2020.12.13.422554

**Authors:** Ying Liu, Bo Ding, Lina Zheng, Ping Xu, Zhiheng Liu, Zhao Chen, Peiyao Wu, Ying Zhao, Qian Pan, Yu Guo, Wei Wang, Wensheng Wei

## Abstract

Increasing evidence shows that promoters and enhancers could be related to 3D chromatin structure, thus affecting cellular functions. Except for functioning through the canonical chromatin loops formed by promoters and enhancers, their roles in maintaining broad chromatin organization have not been well studied. Here, we focused on the active promoters/enhancers (referred to as hotspots) predicted to form many 3D contacts with other active promoters/enhancers, and identified dozens of loci critical for cell survival. While the essentiality of hotspots is not resulted from their association with essential genes, deletion of an essential hotspot could lead to change of broad chromatin organization and expressions of distal genes. We demonstrated that multiple affected genes that are individually non-essential could have synergistic effects to cause cell death.

## INTRODUCTION

In the human genome, promoters and enhancers can be brought to spatial proximity by chromatin loops to form regulatory interactions^1–4^. Promoter-enhancer contacts are often mediated by specific DNA binding proteins that form a complex. Imaging analyses have shown that transcription factors (TFs) and polymerase are not evenly distributed in the nucleus but rather concentrated in certain regions to form spatial clusters^1–4^. Furthermore, recent studies have revealed that transcription elongation can be critical for the chromatin organization^5^. These studies suggested a mutual relationship between promoter/enhancer activity and 3D chromatin structure.

Given these observations, we hypothesize the following: if a promoter or enhancer is positioned in the 3D space such that it is pivotal for maintaining and stabilizing the surrounding 3D chromatin structure, perturbing this element may impact the chromatin organization beyond their interacting loci; namely, perturbation of such promoter or enhancer would significantly alter broad chromatin organization and impact transcription of non-direct target genes. The current studies of promoter/enhancer functions have been largely focused on the direct regulation of gene expression, but their roles in organizing broad chromatin structure, i.e. their “structural importance”, has been largely overlooked.

Here, we aim to investigate this hypothesis and focus on the active promoters/enhancers that likely form many 3D contacts with other active promoters or enhancers, referred to as hotspots hereinafter. Using the previously published algorithm EpiTensor^6^, we identified hotspots based on their covariation of epigenetic marks across cell types, and found that the cancer-specific genetic variations (GVs) have significantly higher chance to reside in hotspot regions. Through a high-throughput CRISPR-Cas9 library screening of hotspots by targeted deletion, we identified dozens of loci critical for cell growth and survival. Importantly, we showed that the cell death was not caused by the dysregulation of their direct target genes, but rather by impacting genes to generate synergistic effect at the systems level that alters broad chromatin organization and gene expression.

## RESULTS

### Small world network formed by the 3D contacts between promoters and enhancers

To identify promoters or enhancers that are likely important for chromatin organization, we set out to identify such elements involved in multiple interactions with other loci in the genome. Because Hi-C data with sufficient resolution to define the interactions between promoters and enhancers are rare, we resorted to computational prediction. We have previously shown that chromatin contacts could be successfully predicted by the covariation of epigenetic modifications in the contacting loci^6^. Briefly, we developed an unsupervised learning method called EpiTensor^6^ that could detect epigenetic covariation patterns between promoter-enhancer, promoter-promoter and enhancer-enhancer pairs at 200-bp resolution using tensor analysis. Such a covariation indicates possible 3D contacts, which can be realized in a cell-type specific manner. Therefore, when considering spatial contacts in a particular cell type or tissue, we focused on those formed between active promoters and enhancers because these contacts are likely to establish functional regulation. We showed that the EpiTensor predictions were highly concordant with the Hi-C, ChIA-PET and eQTL results in different cell types^6^.

Here, we identified active promoters (marked by H3K27ac and H3K4me3) and enhancers (marked by H3K27ac and H3K4me1) in 73 normal and 5 cancer cells/tissues with all 3 marks available in the Roadmap Epigenomics project^7^. The 3D contacts between these *cis*-elements predicted by EpiTensor^6^ were assembled into a regulatory element interaction network (referred to as REIN hereinafter) in each cell/tissue and its topological properties were analyzed using SNAP^8^. The REINs are small world network because their cluster coefficients (the percentage of nodes’ pair connected when they both connected to another node) are similar to an equivalent (same number of nodes and edges) regular lattice network^9^ and their path length (the largest required number of steps between node pairs) is similar to that of the equivalent random network^10^ (**Fig. 1a**). Small world network is robust in that it is resistant to random attack but vulnerable to targeted attack on the high degree nodes^11^. Therefore, we selected the top 10% high degree nodes in REIN, referred to as hotspots thereinafter, for further analysis.

**Figure 1.**
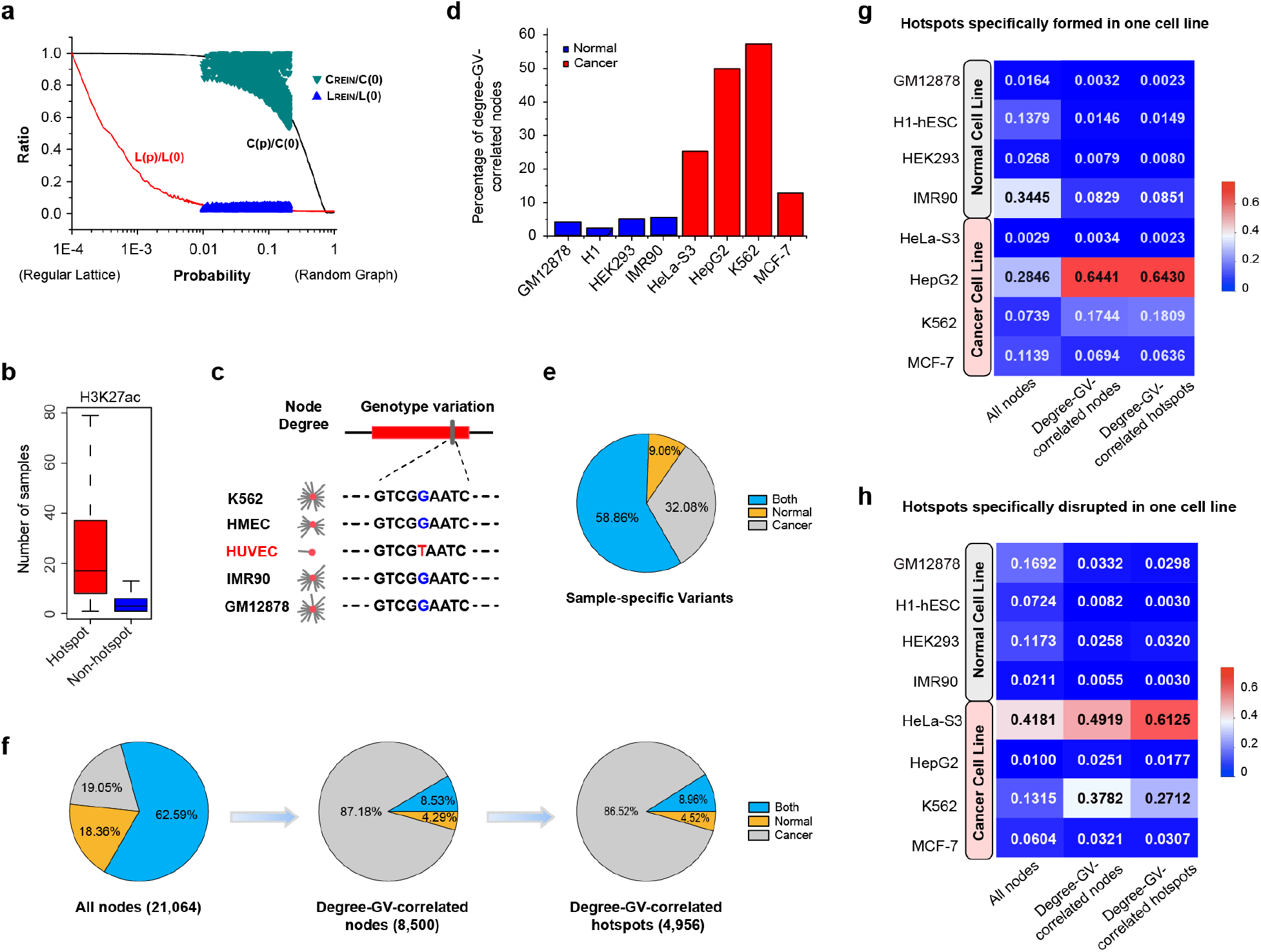
Small world network analysis and mutations effects on 3D contact for hotspot enhancers and promoters. (**a**) The path length and cluster coefficient of REINs compare with equivalent regular lattice networks and equivalent random graph networks. (**b**) Comparison of H3K27ac peaks between hotspot and non-hotspot in 121 cell lines/primary cells/tissues characterized by the NIH Roadmap Epigenetics Project. (**c**) Definition of degree-GV-correlated nodes. In this example, the node has low degree in HUVEC and high degree in other cell lines, which is correlated with the GV profile with a G > T SNP in HUVEC but none in other cell lines. (**d**) The percentage of Degree-GV-correlated-nodes in normal cell lines and cancer cell lines. (**e**) The distribution of GV specificities in samples. Normal, Cancer and Both indicate the GVs with specificities only in normal, cancer and both samples, respectively. (**f**) The distribution of normal or cancer cell-line specificities in All-nodes, Degree-GV-correlated-nodes and Degree-GV-correlated-hotspots. (**g**) The distribution of one-cell-line-hotspot-formation in all-nodes, Degree-GV-correlated-nodes and Degree-GV-correlated-hotspot. (**h**) The distribution of one-cell-line-hotspot-disruption, in all-node, Degree-GV-correlated-nodes and Degree-GV-correlated-hotspot.

### Mutations on hotspot enhancers and promoters could alter 3D contacts

We collected all the genomic loci identified as hotspots in at least one cell/tissue. In total, we found 48,110 regions, and the majority of them are enhancers while 12,754 of them overlap with promoter regions (1-kb around the transcription start sites). Consistent with our previous analysis^6^, these loci tend to be active (overlapping with H3K27ac signals) in more cell types than the non-hotspot loci (**Fig. 1b**). We noticed that the number of interactions a hotspot forms varies significantly across cell types/tissues and on average a locus was identified as a hotspot only in 7 out of 78 cell types/tissues. Particularly, promoter hotspots are shared by more cell types/tissues (on average 17 out of 78) than enhancer hotspots (on average 4 out of 78). Because enhancers are known to be cell type/tissue specific, it is not surprising that potential hotspot enhancers only form many contacts in specific cells/tissues.

Given the importance of high-degree nodes in a small world network, mutations on the hotspot loci may have severe impact on the network structure. To investigate this possibility, we analyzed the loci that are active in both normal and cancer cell lines but with a significant degree change. We first identified nodes with sample specific degrees: using the degree numbers of each node in all 73 normal samples as the background distribution, we identified nodes that are active in a specific sample and whose degree also significantly deviates from the mean. We then determined the sample specific genetic variations (GVs) (referred to single nucleotide variations (SNVs), hereinafter). We collected 1,197,917 GVs in 45 normal and 17 cancer samples (DCC accession number ENCFF105JRY). For each GV in each sample, if its B-allele frequency significantly deviates from the mean in the 45 normal samples, we considered this GV specific to the sample.

The nodes with sample specific degree containing at least one sample specific GV, which are called degree-GV correlated nodes (**Fig. 1c**), are good candidates to investigate the relationship. We first analyzed 4 normal (GM12878, H1, HEK293 and IMR90) and 4 cancer cell lines (HeLaS3, HepG2, K562 and MCF-7), and found that the degree-GV correlated nodes are more frequently observed in cancer cell lines than normal cell lines (**Fig. 1d**). Among 21,064 nodes with specific high/low degree in at least one cell line we found that the majority (62.59%) showed specificities in both cancer and normal cells, 18.36% only in cancer, and 19.05% only in normal cells (**Fig. 1f**). Similarly, among the 629,547 cell-specific GVs, 58.86% showed specificities in both cancer and normal cells, 32.08% only in cancer and 9.06% only in normal cells (**Fig. 1e**). However, the degree-GV-correlated nodes were dominated by cancer-specific ones (87.18%), compared to 8.53% showing specificities in both cancer and normal cells, and 4.29% only in normal cells (**Fig. 1f**). We observed the same trend for degree-GV-correlated hotspots, including 86.52% cancer-specific, 4.52% normal-specific, and 8.96% in both (**Fig. 1f**). In summary, the dominant majority of degree-GV-correlated nodes appear in cancer cells.

We further examined two groups of nodes in the 8 distinct cell lines, one group having significant higher degree in one cell line than in other cell lines that indicates cell-type specific contact formation (one-cell-type-specific nodes), and another having significant lower degree in one cell line than in the others that indicates cell-type specific contact disruption (seven-cell-type-specific nodes). The percentages of HepG2-specific nodes and K562-specific nodes in one-cell-type-specific group (cell-type specific contact formation) are 28.5% and 7.4% in all nodes, 64.4% and 17.4% in degree-GV-correlated-nodes and 64.3% and 18.1% in degree-GV-correlated-hotspots, respectively (**Fig. 1g**). Similarly, the percentages of HeLa-S3-specific nodes and K562-specific nodes in seven-cell-type-specific group (cell-type specific interaction disruption) are 41.8% and 13.2% in all nodes, 49.2% and 37.8% in degree-GV-correlated-nodes and 61.3% and 27.1% in degree-GV-correlated-hotspots, respectively (**Fig. 1h**). Taken together, our analyses suggested that cancer-specific GVs are highly correlated with the node-degree change that alters REIN.

### CRISPR/Cas9 library screening identified hotspots essential for cell growth and survival

To further investigate the function of hotspots, 751 hotspots identified as enhancers were randomly selected for targeted deletion to analyze their impact on cell growth (**Supplementary Table 1**). These hotspots do not overlap with coding regions of any protein-coding gene or non-coding RNA. In total, 14,399 paired-gRNAs (pgRNAs) were designed to delete these loci, including positive control pgRNAs targeting ribosomal genes, negative control pgRNAs targeting *AAVS1* locus and non-targeting pgRNAs (**Supplementary Table 2**). Through lentivirus infection at a low MOI, the pgRNA library was transduced into K562 cells stably expressing Cas9 protein. The pgRNA-infected samples were FACS-sorted 3 days post infection, serving as the control group, then were continuously cultured for 30 days to obtain the experimental group. The library cells from the control and experimental group were sequenced to determine the abundance of each pgRNA (**Fig. 2a**). The read distribution of pgRNAs showed high correlation between the two biologically independent replicates for all groups (**Supplementary Fig. 1a-c** and **Supplementary Table 3**), indicating high reproducibility.

**Figure 2.**
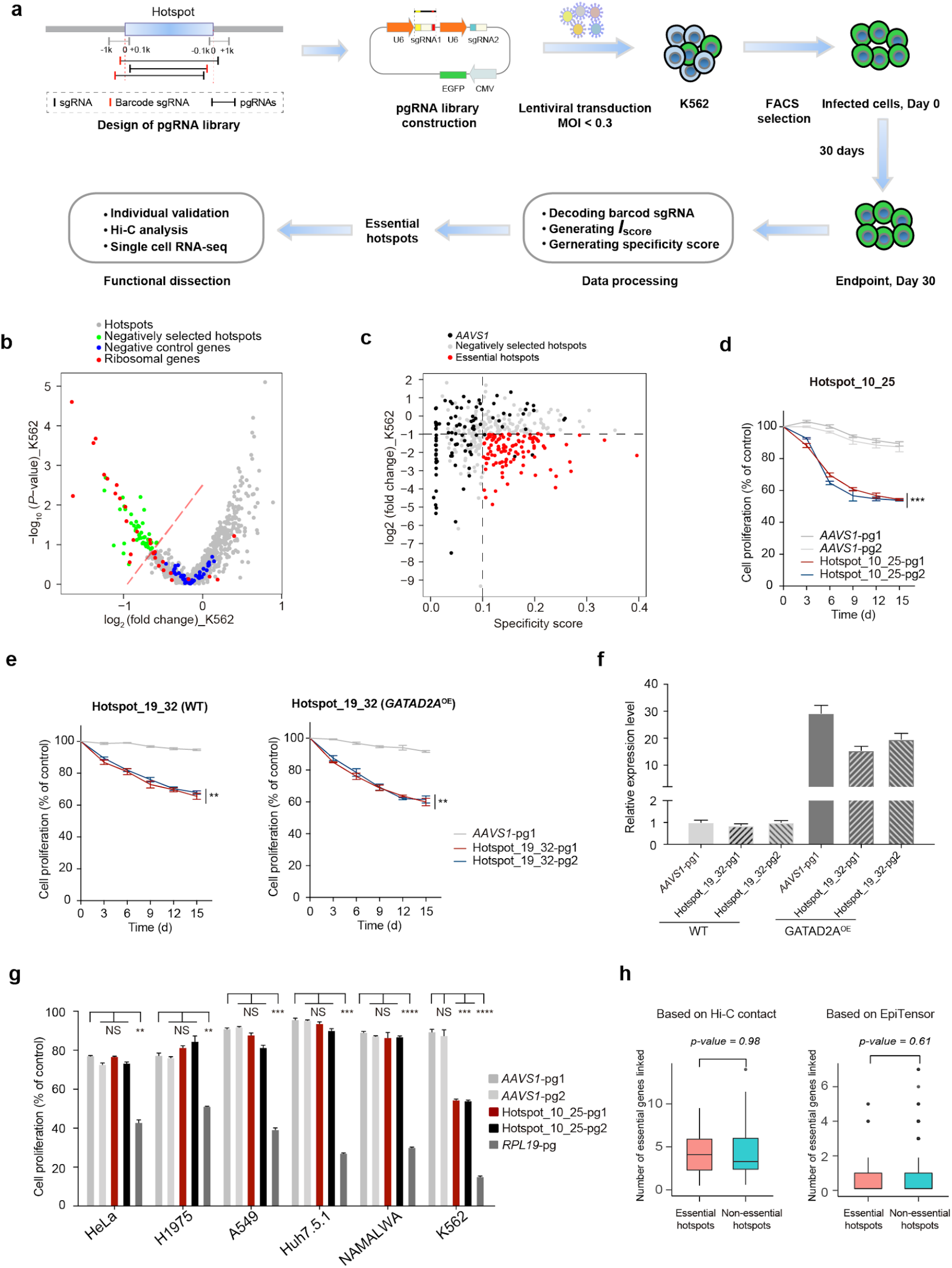
Identification of essential hotspots for cell growth and proliferation in the K562 cell line through pgRNA-deletion-based CRISPR screening. (**a**) The Schematic of the pgRNA library design, cloning and functional screening of selected hotspot loci. (**b**) Volcano plot of the fold change and *p-value* of hotspots in the K562 cell line. Negative control genes were generated by randomly sampling 20 *AAVS1*-targeting pgRNAs with replacement per gene, and ribosomal genes served as positive control in the screening. Dotted red line represents *Iscore* = −2.5. (**c**) Selection of candidate essential hotspots by the fold change and specificity score of each pgRNA. These essential hits were selected under the threshold of specificity score > 0.1, log2 (fold change) < −1. (**d**) Validation of top-ranked essential hotspot in K562 cell by cell proliferation assay. *AAVS1*-pg1 and *AAVS1*-pg2 are pgRNAs targeting *AAVS1* locus as negative controls. Asterisk (*) represents *p-value* compared with pgRNAs targeting *AAVS1*-pg1 at Day 15, calculated by two-tailed Student’s *t*-test and adjusted for multiple comparisons by Benjamini–Hochberg procedure. Data are presented as the mean ± s.d. (n = 3 biologically independent samples). * *p* < 0.05; ** *p* < 0.01; *** *p* < 0.001; **** *p* < 0.0001; NS, not significant. The pgRNAs for individual validation of each hotspot are listed in **Supplementary Table 5**. (**e**) Validation of essential hotspot overlapped with the intronic region of an essential gene in K562 cell by cell proliferation assay. Left: WT K562 cells infected with pgRNAs targeting hotspot_19_32. Right: *GATAD2A-*overexpressed K562 cells infected with pgRNAs targeting hotspot_19_32. (**f**) The expression levels of *GATAD2A* of WT and *GATAD2A-*overexpressed K562 cells infected pgRNAs targeting *AAVS1* or hotspot_19_32. (**g**) Validation of hotspot_19_32 in multiple cancer cell lines including A549, H1975, HeLa, Huh7.5.1 and NAMALWA cell lines. Asterisk (*) represents *p-value* compared with pgRNAs targeting *AAVS1*-pg1 at Day 15, calculated by two-tailed Student’s *t*-test and adjusted by Bonferroni correction accounting for multiple testings. * *p* < 0.05; ** *p* < 0.01; *** *p* < 0,001; **** *p* < 0.0001; NS, not significant. (**h**) No significant difference between the numbers of essential genes contacting the essential and non-essential hotspots from Hi-C or EpiTensor in K562. The pgRNAs used above were listed in **Supplementary Table 5**.

Compared with the control group, pgRNAs in Day-30 experimental cells targeting ribosomal genes, hotspots were both decreased more than those targeting *AAVS1* locus and non-targeting pgRNAs. We calculated the *I*_*score*_ for each hotspot (**Online Methods** and **Supplementary Table 4**), and 49 hotspots with scores ≤ −2.5 were considered to significantly affect cell fitness upon deletion (**Fig. 2b**). To avoid the cellular toxicity caused by off-target effects^12–15^, we assessed the specificities of sgRNAs with 2 or 3 mismatches to off-target loci using GuideScan specificity score^16^. Because *AAVS1*-targeting pgRNAs with specificity score ≤ 0.1 could cause dropout effect in K562 (Fig. 2c and Supplementary Fig. 1d), we only kept pgRNAs with specificity score > 0.1 and log_2_(fold change) < −1 for subsequent analysis. Furthermore, hotspots with copy number amplification were also removed to avoid cell death caused by multiple cleavages^17^. Using such a stringent criteria, we identified 43 hotspots essential for cell viability of K562 cells (**Fig. 2c**).

Based on the ranking of *I*_*score*_, 7 top-ranked hotspots in K562 were chosen for individual validation by cell proliferation assay. All of them do not overlap with any promoter, protein-coding gene or non-coding RNA. Three or two pgRNAs with high targeting specificity were constructed separately for each hotspot, and cell proliferation assay was performed as previously reported^18^. We found that deletion of these hotspots led to significant cell death or cell growth inhibition (**Fig. 2d** and **Supplementary Fig. 2a**), which was consistent with the screening results, indicating that these hotspots played critical roles in cell fitness.

For hotspots located in promoters or intronic regions, their deletion may affect the expression of the host genes. We investigated whether the cell death caused by hotspot deletion is a trivial effect of inhibiting expression of essential genes. For hotspot_19_32 located in the intron of an essential gene *GATAD2A*, we chose 2 highly-specific pgRNAs to respectively delete this locus in K562 cells, and observed significant cell growth inhibition (**Fig. 2e**). Importantly, we found that overexpression of *GATAD2A* did not rescue the cell death caused by hotspot deletion (**Fig. 2e**), indicating that the hotspot deletion has a profound impact on cell growth. Through detecting the expression level of the gene *GATAD2A* in each condition by real-time qPCR, we confirmed that the gene was successfully overexpressed in K562 cells and the cell death caused by hotspot deletion was not rescued by *GATAD2A* overexpression (**Fig. 2f**). Similar result was obtained for hotspot_1_36, which is located about 3-kb upstream of the transcriptional start site of an essential gene *SLC2A1*. We performed the cell proliferation assay using 2 pgRNAs in the wild type K562 cell and K562 cell stably expressing *SLC2A1*. Similar level of cell fitness influence was observed in both conditions for each pgRNA deletion, and real-time qPCR further confirmed that the growth phenotype was not due to affecting the expression level of *SLC2A1* (**Supplementary Fig. 2b-c**).

To further assess the essentialities of the identified K562-essential hotspots in other cancer cell lines, we chose hotspot_10_25, which showed significant growth defect in K562 if deleted, for parallel validations in HeLa (cervical cancer cells), H1975 (non-small cell lung cancer cells), A549 (non-small cell lung cancer cells) and NAMALWA (Burkitt’s lymphoma) cells. Surprisingly, compared with the negative control *AAVS1*-targeting pgRNAs, hotspot_10_25 showed no significant effect in any of the five tested cell lines, indicating its role in K562 cells is cell type specific (**Fig. 2g** and **Supplementary Fig. 2d**).

### Essentiality of hotspots is not resulted from association with essential genes

To analyze the mechanism of how these identified essential hotspots exerts their functional roles, we first examined whether direct interaction with essential genes determines the essentialities of these hotspots. We collected the essential genes according to the CRISPRi-based screen^19^ and identified all possible spatial contacts they formed that were detected by Hi-C (*p*-value ≤ 0.05) in the wild type K562 cells^20^. There is no distinction between essential and non-essential hotspots regarding their association with essential genes (the *Wilcoxon* Rank-Sum test *p*-value = 0.98 indicating no significant difference) (**Fig. 2h**). We also performed the same comparison using the spatial contacts predicted by EpiTensor and obtained the same observation (the *Wilcoxon* Rank-Sum test *p*-value = 0.61) (**Fig. 2h**). According to the above analysis, the essentiality of hotspots is not determined by their direct contact with essential genes.

### Deleting essential hotspots affects broad chromatin organization

We next investigated whether deleting hotspot affects the chromatin organization. We selected hotspot_10_25 for further analysis, which showed unique essentiality in K562 (**Fig. 2d, g** and **Supplementary Fig. 2d**) and does not interact with any essential protein-coding gene. We first performed whole genome sequencing (WGS) to confirm that there was no off-target issue. The validated pgRNA hotspot_10_25-pg2 was chosen (**Fig. 2d**), and the WGS library was generated 8 days post pgRNA infection in K562 cells. Compared to the hg19 human genome, we successfully identified 4.1 million germline mutations in hotspot_10_25-deleted K562 cells, 86.2% consistent with the published K562 wildtype WGS data. The high percentage of the germline mutation discovery rate indicated good quality of the library. We used Cas-OFFinder to identify 746 potential off-target loci with loose cutoffs (base mismatch ≤ 4, bulge ≤ 2) to avoid missing any possible off-target loci. We manually examined the putative off-target loci with the indels detected from the edited cells that differed from the wild type (Supplementary Table 6). Except for the significant indels found in the two on-target loci, there were no hits on any of the putative off-target loci. These analyses confirmed that cell proliferation was not resulted from off-target effect.

We subsequently performed Hi-C analysis on the edited K562 cells and compared that with the wild type^20^ (see Methods). No significant difference was observed through visual inspection of the Hi-C contact maps at 100-kb, 25-kb and 10-kb resolution (**Fig. 3a**). For a quantitatively assessment, we evaluated the global characteristics of the contact networks constructed from the Hi-C data, i.e. each node in the network is a 5 kb-long locus and edge is a significant contact detected by Hi-C (5-kb VC normalized reads with the cutoff of *p*-value ≤ 0.05). Because the Hi-C data had a limited number of inter-chromosomal contact reads, we focused on intra-chromosomal contacts and constructed a network for each individual chromosome. We identified modules (i.e. densely connected nodes) for each chromosome using the SNAP software^21^ with the Clauset-Newman-Moore algorithm^22^. Hotspot deletion had no significant impact on the chromosome-wide modularity (i.e. the difference between the fraction of edges observed within a group of nodes and the expected value in random network) and effective diameter (i.e. the path length such that 90 percent of node pairs are a smaller or equal distance apart) as indicated by a strong correlation R = 0.88 for modularity scores and R = 0.74 for effective diameter of the wild type K562 and hotspot-deleted networks in 23 chromosomes (**Supplementary Fig. 3**), which is consistent with the visual inspection of the Hi-C contact maps.

**Figure 3.**
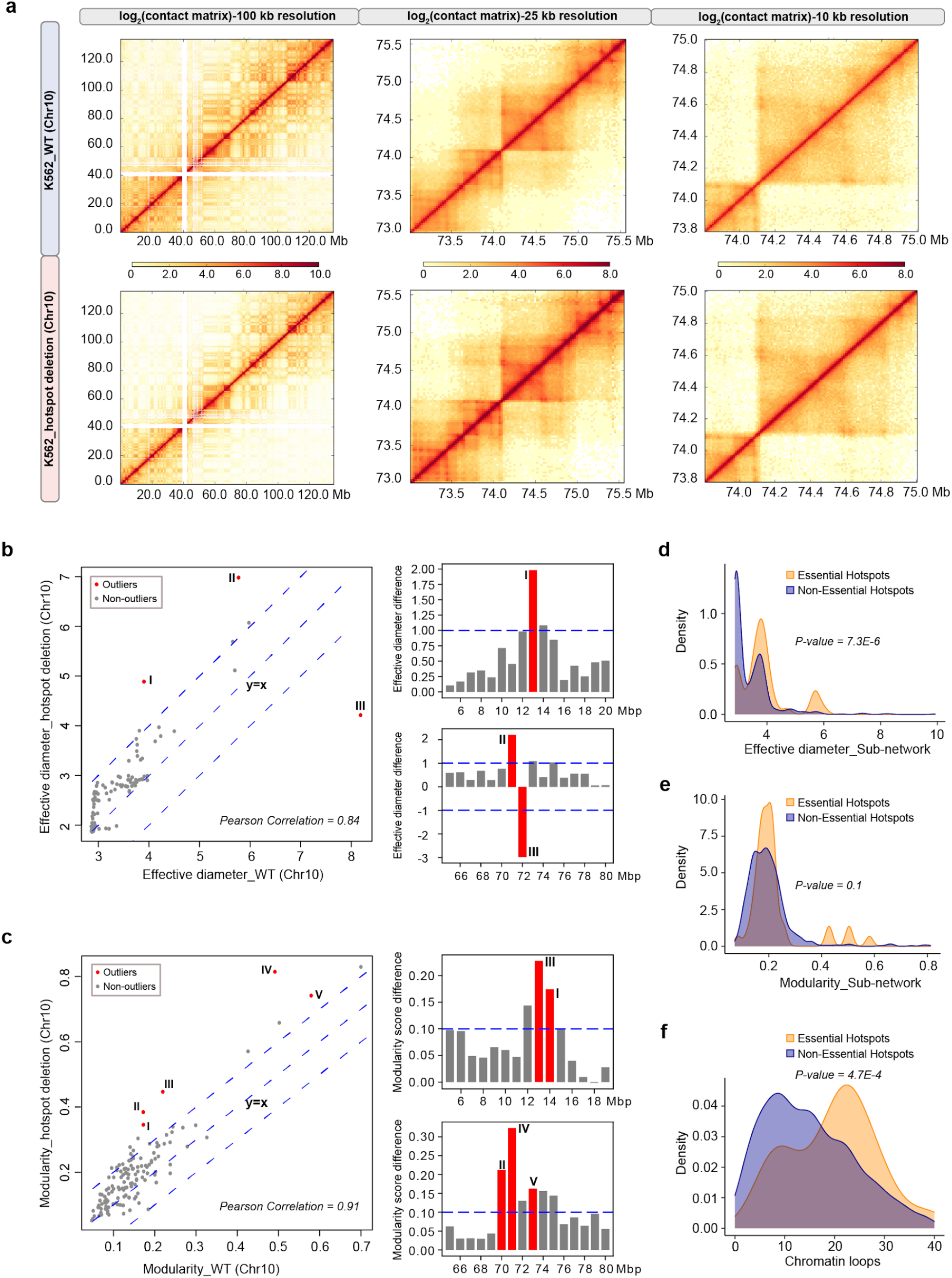
Deletion of essential hotspot impacts broad chromatin structure. (**a**) Hi-C contact maps of the entire chr10 at 100-kb resolution (left), chr10: 73 Mb-75.5 Mb at 25-kb resolution (middle) and chr10: 73.8 Mb-75.0 Mb at 10-kbp resolution (right). (**b-c**) The effective diameter (**b**) and modularity (**c**) before and after hotspot deletion in the sliding 5-Mbp sub-networks in chr10 (left). The outliers are labeled and their genomic locations are shown on the right. (**d-f**) The distributions of the effective diameter (**d**), modularity score (**e**) and chromatin loops (**f**) in the 5-Mbp regions around the essential hotspots and non-essential hotspots.

Without finding chromosome-wide difference of chromatin organization upon hotspot deletion, we investigated whether deleting a hotspot could affect the neighboring regions in the spatial proximity. Using a sliding window of 1-Mb, we identified all Hi-C contacts (5-kb resolution with p-value ≤ 0.05) that each 1-Mb segment forms with its flanking 2-Mb regions at both sides in the linear genome and these contacts were assembled into a sub-network. The modularity score and effective diameter were computed in each of these sub-networks for the wild type and hotspot-deleted K562 cells. These two metrics showed high chromosome-wide correlation in chr10 before and after hotspot deletion, with a Pearson correlation of 0.84 and 0.91 for effective diameter and modularity, respectively (Fig. 3b-c). Noticeably, significant changes were observed surrounding the hotspot for the network characteristics, effective diameter (chr10: 69 Mb-75 Mb) and modularity (chr10: 68 Mb-76 Mb) (**Fig. 3b-c**). Some genomic regions interacting with hotspot neighbor regions were also affected such as chr10: 11 Mb-17 Mb showing significant change of modularity (**Fig. 3c** and **Supplementary Fig. 4**). These analyses showed that hotspot deletion resulted in a broad chromatin structure alteration.

**Figure 4.**
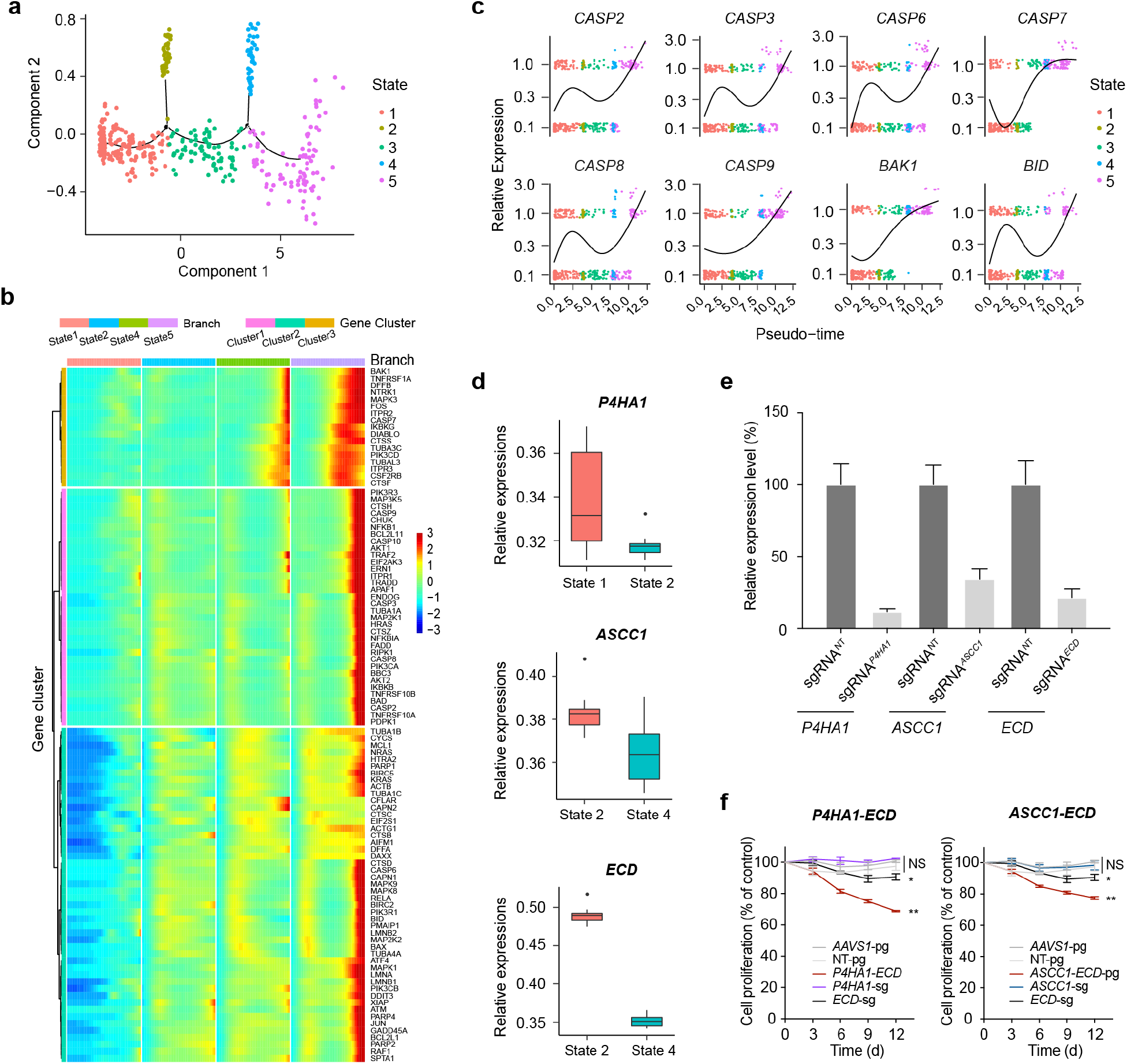
Synergistic change of gene expression after hotspot deletion. (**a**) Pseudotime clusters of hotspot_10_25-deleted and wild type K562 cells based on apoptosis gene expressions. (**b**) Relative expressions of typical apoptosis genes (*CASP2, CASP3, CASP6, CASP7, CASP8, CASP9, BAK1, BID*) under different pseudotime plotted using the Monocle package. Dots in different colors represent different cell states. (**c**) Global analysis of the expression levels of 99 KEGG apoptosis genes in state 1, 2, 4 and 5. Genes were clustered into 3 groups. (**d**) The Relative expression levels of three representative down-regulated genes in different states from single cell RNA-seq. (**e**) The knockdown efficiency of the indicated sgRNAs targeting each down-regulated gene in K562 cells stably expressing KRAB-dCas9. The expression level of each gene was detected by real-time qPCR. sgRNA^NT^ represents the non-targeting sgRNA serving as the negative control. (**f**) Validation of the synergistic effects of two gene pairs on K562 cell fitness by cell proliferation assay. Asterisk (*) represents *p-value* compared with pgRNAs targeting *AAVS1*-pg at Day 12, calculated by two-tailed Student’s *t*-test and adjusted for multiple comparisons by Benjamini– Hochberg procedure. * *p* < 0.05; ** *p* < 0.01; NS, not significant. The sgRNAs, pgRNAs and primers were listed in **Supplementary Table 5**.

### Essential hotspots tend to reside in dense chromatin structure

If the essential hotspots are critical for maintaining the chromatin structure in the spatial neighborhood, it is likely that the 3D contacts around them are dense. Therefore, we compared the sub-network effective diameters, modularity and chromatin loops in the 5-Mb regions centered at the essential and nonessential hotspots in the wild type K562. We found that essential hotspots were surrounded with higher effective diameters (*Wilcoxon* Rank-Sum test, *p*-value = 7.3E-6), higher modularities (*p*-value = 0.1) and higher loop densities (*p*-value = 4.7E-4) than the non-essential ones (**Fig. 3d-f**). In fact, using these three metrics in the wild type K562 cells, a random forest classification model could distinguish essential from non-essential hotspots with an AUC of 0.73 in 10-fold cross validations. This result resonates with the above observations and suggests that hotspots are pivotal for stabilizing the dense chromatin contacts in the spatial neighborhood.

### Hotspot deletion synergistically affects gene expression to cause cell death

To further investigate the mechanisms of how hotspot deletion leads to cell death or cell growth inhibition, we performed single cell RNA-seq using Drop-seq^23^ to analyze the gene expression change upon hotspot_10_25 deletion. We transduced the individually validated pgRNA hotspot_10_25-pg2 (**Fig. 2d**) targeting this essential hotspot into K562 cells, among which 482 single cells passed the quality control. We also included the bulk RNA-seq data of wild type and *AAVS1*-deleted cells as controls. All the data were normalized together to make them comparable (**Online Methods**). As deletion of this hotspot has an impact on cell viability or cell growth, we focused on the apoptosis pathways to confirm their activation. We selected 99 apoptosis genes documented in the KEGG database and clustered the cells into five states by trajectory branching and pseudotime analysis using Monocle (**Fig. 4a**)^24^. The wild type and *AAVS1*-deleted cells were located in state 1, suggesting that cells in this state resemble the wildtype. The apoptosis genes fell into three groups, with distinct expression patterns along the pseudotime, for example, *CASP2, CASP8, CASP9*, and *CASP10* in cluster 1, *CASP6* in cluster 2, and *CASP7* in cluster 3 (**Fig. 4b**). Overall, there was a trend of increasing gene expression from state 2 to state 5, such as *CASP2, CASP3, CASP6, CASP7, CASP8, CASP9, BAK1* and *BID* (**Fig. 4c**). The transcriptomic analysis showed that apoptosis pathways are activated upon hotspot deletion.

To investigate the impact of hotspot deletion on the spatial neighborhood, we analyzed the genes whose promoters were predicted to interact with the essential hotspot_10_25 by EpiTensor. Among the 14 genes located within the same TAD of hotspot_10_25, 4 of them were significantly down-regulated in the progress from state 1 to 5, including *P4HA1* (down-expressed from state 1 to 2, **Fig. 4d**), *DNAJB12, ASCC1* and *ECD* (down-expressed from state 2 to 4, **Fig. 4d** and **Supplementary Fig. 5a**). Through individually knocking down each gene by CRISPR interference (**Fig. 4e** and **Supplementary Fig. 5b**), only *ECD* showed a weak impact on cell growth and all the other genes showed no detectable effects (**Fig. 4f** and **Supplementary Fig. 5c**). As the hotspot interacted with multiple genes, we investigated whether knocking down a pair of genes would have the synergistic effect on cell growth. Applying the CRISPRi strategy, we knocked down 6 pairs of genes (**Supplementary Table 5**) using paired-gRNAs in K562 cells, respectively. We found that simultaneous knockdown of *P4HA1-ECD* and *ASCC1*-*ECD* showed much more significant impact on cell growth (**Fig. 4f**). These results indicated that disrupting hotspot_10_25 could affect the expression levels of multiple interacting genes, and their synergistic effect led to cell death or cell growth inhibition. Note that we were limited to examine pairs of genes but hotspot deletion can affect multiple genes together with more significant synergic effects.

## DISCUSSION

In this study, we have revealed the understudied “structural importance” of regulatory elements especially enhancers. We observed that genetic variations in cancer cells are strongly correlated with alteration of 3D contact degrees in the hotspots. Furthermore, we showed that deleting a hotspot enhancer could have a broad impact on chromatin structure that is beyond chromatin looping and affect genes not directly interacting with the enhancer. The affected genes could have synergistic effects to inhibit cell proliferation and cause cell death, although each individual one is not critical for cell survival. These observations suggested that at least some enhancers have pivotal roles of stabilizing proper chromatin organization in a broad scale that has not been fully appreciated.

## METHODS

Methods, including statements of data availability and any associated accession codes and references, are available in the online version of the paper.

## ACKNOWLEDGEMENTS

We acknowledge the staff of the BIOPIC High-throughput Sequencing Center (Peking University) for their assistance in next-generation sequencing analysis, the National Center for Protein Sciences (Beijing) at Peking University for assistance with Fluorescence-activated cell analysis and sorting, and Dr. H. L., Ms. L. D. for technical help. We acknowledge Dr. Ying Yu (Peking University) for the assistance in preparing the NGS library. We acknowledge the staff of UC San Diego IGM Genomics Center for sequencing service and UC San Diego Human Embryonic Stem Cell Core Facility for cell sorting service. We acknowledge Ms. Jia Xu (UC San Diego) for the assistance in preparing a single cell RNA-seq library. This project was partially supported by CIRM (RB5-07012) and NIH (R01HG009626) (to Wei Wang), and was supported by funds from the National Science Foundation of China (NSFC31930016), Beijing Municipal Science & Technology Commission (Z181100001318009), the Beijing Advanced Innovation Center for Genomics at Peking University, and the Peking-Tsinghua Center for Life Sciences (to Wensheng Wei).

## AUTHOR CONTRIBUTIONS

W.Wei and W.Wang conceived and supervised the project. W.Wei, W.Wang, Y.L. and B.D. designed the experiments. B.D and L.Z. constructed network analysis, identified and characterized hotspot regions. Y.G. designed the pgRNA library for hotspot screening. Y.L. and P.X. performed the pgRNA library construction and screening. Y.L. performed the experiments including individual validation of candidate hotspots in multiple cell lines, whole genome sequencing (WGS), bulk RNA-seq and examination of the synergistic effects with the help of P.X. and Q.P.. Z.L. performed the bioinformatics analysis of the screening data and designed the pgRNAs used for individual validation. P.W and Z.C. performed the Hi-C experiments on hotspot-deleted K562 cells. P.W. and Y.Z. performed single cell RNA-seq on hotspot-deleted K562 cells. L.Z. performed the bioinformatics analysis of the single cell RNA-seq and Hi-C data. Y.L., B.D., L.Z., W.Wang and W.Wei wrote the manuscript with contributions from all other authors.

## COMPETING FINANCIAL INTERESTS

The authors declare no competing financial interests.

## ONLINE METHODS

### Cell culture

K562, H1975 and NAMALWA cells were cultured in RPMI 1640 medium (Gibco), 293T, HeLa, A549 and Huh7.5.1 cells were cultured in Dulbecco’s modified Eagle’s medium (DMEM, Gibco). All cells were supplemented with 10% fetal bovine serum (FBS, Biological Industries) with 1% penicillin/streptomycin, cultured with 5% CO_2_ in 37°C.

### Design and construction of the CRISPR-Cas9 pgRNA library

To explore the cellular function of hotspots, we selected 751 hotspots identified in the K562 cell line. For each hotspot, sgRNAs targeted 100-bp inside regions and 1-kb outside regions flanking the two boundaries of hotspot loci, respectively. If there were not enough sgRNAs satisfying the following design rule, sgRNAs were searched among the 5-kb outside regions flanking each boundary. All the PAM motifs in targeting regions were scanned to identify available sgRNA targeting sites. All the selected sgRNAs are located in noncoding regions and satisfy all the following conditions: (1) The targeting sequence is unique for the intended locus; (2) The sgRNA contains at least 2 mismatches to any other locus in the human genome; (3) The GC content of the sgRNA ranges from 20% to 80%. We enumerated all possible pgRNAs from the selected sgRNAs, and then retained those satisfying these conditions: (1) The two sgRNAs respectively targeted 100-bp inside regions or 1-kb (or 5-kb) outside regions flanking each hotspot boundary; (2) The deletion regions should not overlap with any promoter or exonic region of protein-coding genes; (3) The sgRNA targeting sites are at least 30 bp away from the exon-intron boundary of protein-coding genes. The gRNA pairs were designed with one unique gRNA serving as decoding barcode and up to 20 pgRNAs were designed for each locus.

Finally, 14,399 pairs of gRNAs targeting 751 hotspots were generated for the hotspot deletion library, together with 473 pgRNAs targeting the promoter regions (5-kb upstream of the transcription start site) and the first exon of 29 ribosomal genes serving as positive controls, as well as 100 pgRNAs targeting *AAVS1* locus and 100 non-targeting pgRNAs from previous library^25^ serving as negative controls. According to the two-step cloning method^25^, the 128-nt oligonucleotides containing pgRNA coding sequences were synthesized (Agilent Technologies, Inc.), cloned into the lentiviral expression vector harboring an EGFP selection marker (with a minimum representation of 150 transformed colonies per pgRNA in each cloning step) and further packaged as previously described^25^.

### CRISPR-Cas9 pgRNA library screening

To ensure the infection at 1,000-1,500 cells per pgRNA with an MOI of < 0.3, K562 cells stably expressing Cas9 were seeded in duplicate in the T-175 flasks (Corning). 24 hours later, each replicate was infected by the pgRNA library lentiviruses supplemented with 8 μg/ml of polybrene. 72 h post infection, EGFP^+^ cells were sorted by FACS (Day-0 control group). For each replicate, the initial EGFP^+^ pool (1500-fold coverage) was isolated for DNA extraction, and the same number of cells as the experimental group was maintained at a minimum coverage of 1,500 cells per pgRNA at each passage for 30 days. Then cells of each condition with 1500x library coverage were respectively subjected to genomic DNA extraction, PCR amplification of sgRNA-coding sequence and high-throughput sequencing analysis (Illumina HiSeq2500 platform) as previously described^25^.

### Identification of functional hotspots involved in cell growth and proliferation

The raw pgRNA counts were extracted from paired-end sequencing FASTQ files by bash script based on AWK. Since the low reads in control groups affect the analysis result confidence, the pgRNAs with raw reads less than 5 were eliminated from the following analysis. The total counts were further normalized to adjust the sequence depth of each replicate in both control groups and experiment groups. To further filter noisy pgRNAs, we removed pgRNAs whose quantile difference of two replicates was in either 3% tail of the distribution, 100 negative control genes were generated by randomly sampling 20 *AAVS1*-targeting pgRNAs with replacement. Then, log_2_ (fold change) (log_2_FC) between experimental and control groups were calculated for each pgRNA. Two features for each set of hotspot were calculated: 1) the mean log_2_FC of all pgRNA in the set, denoted by FC_hotspot_; 2) –log_10_*P*_value_ of two-sided Mann Whitney U test of all pgRNA in the set comparing with pgRNAs targeting *AAVS1* locus, denoted by *P*_*hotspot*_. And in order to consider both the fold change and P value, we defined a screen score for the hotspots:

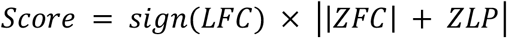

where ZLFC is the Z scaled log_2_FC, and ZLP is the Z scaled –log_10_P_value_. The hotspots with scores of less than −2.5 were identified as essential hotspots.

To further avoid the potential issue of cellular toxicity generated from multiple cleavages by some pgRNAs, we retrived the GuideScan specificity score to evaluate each sgRNA^16^. Each pgRNA was then assigned with half of the harmonic mean of the two sgRNAs’ specificity scores as the pgRNA’s specificity score, which is calculated as follow:

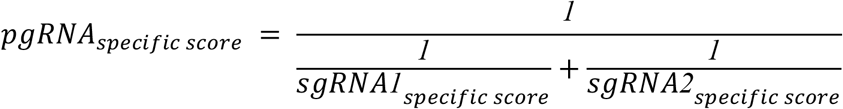

From the identified essential hotspots through the above analysis, those targeting pgRNAs were further selected, whose specificity score is > 0.1 and log_2_ (fold change) is < −1. To further avoid the copy number effects on drop-out screening, the copy number of each hotspot locus in K562 cell line was analyzed based on the ENCODE consortium copy number data (https://www.encodeproject.org/files/ENCFF486MJU/). After filtering hotspot loci with copy number amplification, the remaining hits were regarded as essential hotspots.

### Individual validation of functional hotspots by cell proliferation assay

For each candidate hotspot without immediate overlap with the promoter or gene body of protein-coding genes, two or three pgRNAs were used for the individual validation, which were selected from the library that were consistently depleted or newly designed. To ensure the targeting specificity of all the selected pgRNAs, we required that their specificity scores are all more than 0.15, and the score of at least one pgRNA for each hotspot is more than 0.2. For the newly-designed pgRNA, to ensure the cleavage efficiency, we further required that they don’t include ≥ 4-bp homopolymer stretch and their GC contents are between 0.4 and 0.7. We further ensured each sgRNA targeting site is 400-bp inside and 1-kb outside the two boundaries of hotspot loci, respectively. All the pgRNAs targeting each hotspot locus to be validated were individually cloned into a lentiviral expression vector containing an EGFP selection marker. The cell proliferation assay was performed as previously described^25^. The experiments last for 15 days after the first FACS analysis and at least 100,00 cells were analysed.

For the hotspots overlapping with the promoter or within the intron of possible essential protein-coding genes, three pgRNAs were selected for the subsequent validation. The cDNA of each neighboring coding gene was cloned into lentiviral backbone containing puro selection marker, and individually transduced into K562 cells. Three days post virus infection, the cells with candidate gene overexpression were enriched by puro treatment, then the corresponding pgRNAs targeting the neighboring hotspot were respectively transduced into these cells, as well as the wild type K562 control cells. The cell proliferation assay was performed as described above.

### Hi-C library preparation and data analysis

The pgRNA Hotspot_10_25-pg2 was delivered into K562 cells through lentiviral infection at an MOI of < 1. The EGFP positive cells were collected by FACS sorting at Day 9 post infection, and the sorted cells were allowed to recover in normal cell culture condition for 2 hr before proceeding to Hi-C library. One million cells were used for each Hi-C library preparation using Arima-HiC kit (Arima Genomics, San Diego) following the manufacturer’s instruction. Hi-C libraries were sequenced using the Illumina NovaSeq platform.

An in-house pipeline Juicer^26^ was implemented to process the Hi-C data. Briefly, Hi-C contact reads were aligned to hg19 (GRCh37). All the reads with MAPQ < 30 were further trimmed. The output bam files were transformed into each 10-kb, 25-kb and 100-kb resolution contact matrix, respectively. Then the vanilla coverage (VC) method^27^ was applied on the Hi-C raw reads. Between the expected VC-normalized reads and the observed VC-normalized reads, we fit a *Poisson* Distribution fitting. The normalized contacts were considered as significant if the *p*-value is ≤ 0.05. HiCExplorer^28,29^ and HiCPlotter^30^ were utilized to visualize the processed Hi-C data. HiCCUPS software (https://github.com/aidenlab/juicer/wiki/HiCCUPS) was utilized to call the loops at 10-kb resolution in both the wildtype and hotspot-deleted K562 cells. All the other parameters were set to the default.

### Evaluation of the potential off-target effects by CRISPR-Cas9 system through whole genome sequencing (WGS)

The K562 cells were infected with the validated pgRNA hotspot_10_25-pg2 at an MOI of < 1. Eight days post lentiviral infection, the pgRNA-infected cells were sorted by FACS, and were further subjected to genomic DNA extraction. The whole genome sequencing (WGS) library was prepared following the manufacturer’s instruction, and was sequenced using the Illumina HiSeq 4000 platform. Using the WGS data, we evaluated the potential off-target effects after targeted deletion of hotspot_10_25.

We downloaded the K562 (wild type) WGS data as controls from ENCODE with accession code ENCFF313MGL, ENCFF004THU, ENCFF506TKC and ENCFF066GQD. A strict off-target evaluation was conducted according to the whole genome sequencing approach^31,32^. The putative off-target sites for hotspot_10_25 were generated by Cas-OFFinder in the hg19 genome^31^. In order to avoid any plausible off-target loci, we considered two scenario: 1) base mismatches at most 4 without any bulge mismatch (mismatch ≤ 4, bulge = 0), 2) base mismatches at most 2 with bulge mismatches at most 2 (mismatch ≤ 2, bulge ≤ 2). In total, we examined 746 potential off-target loci. In order to detect the candidate mutations and indels in the edited cells, variant calling was performed as described on GATK Best Practices (https://gatk.broadinstitute.org/hc/en-us). Briefly, reads were aligned to the human reference genome (hg19) using BWA-0.7.17. Duplicated reads were then removed by GATK4 tools MarkDuplicatesSpark (https://gatk.broadinstitute.org/hc/en-us/articles/360037224932-MarkDuplicatesSpark). Lastly, the reads were processed via base quality score recalibration by GATK4 tools. Germline mutations (compared to the hg19 reference genome) were called in both wild type K562 and pgRNA-infected K562 cells by GTAK HaplotypeCaller (version 4.1.4.1). with default parameters. SNVs and Indels in pgRNA-infected K562 compared to the wild type K562 were called by GATK Mutect2 (version 4.1.4.1) with default parameters. Such SNVs and Indels were used to compare with generated putative off-target loci.

For further confirmation, we applied BCFTOOLS suite (version 1.9, http://www.htslib.org/doc/bcftools.html) to call variants. BCFTOOLS mpileup and call commands with default settings were used to generate raw variants. Then, variants with “%QUAL<30 ∥ DP < 30” were marked as low quality variants by BCFTOOLS filter command and filtered out, as well as the homozygous variants with feature “GT=1/1”. We also used BCFTOOLS isec command with parameter “-n −1 -c all” to filter the Mills and 1000G golden standard indels obtained from GATK resource bundle (https://gatk.broadinstitute.org/hc/en-us/articles/360036212652-Resource-Bundle). The putative off-target sites generated by Cas-offinder were checked with the variants called by above BCFTOOLS pipelines, and there were no overlaps found.

### Bulk RNA-seq and data analysis

#### Bulk RNA-seq library preparation

K562 cells were transduced with the pgRNA *AAVS1*-pg1 by lentiviral infection at an MOI of < 1, then 2×10^6^ pgRNA-infected K562 cells were FACS-sorted 8 days post infection. The total RNA was extracted using RNeasy Mini Kit (QIAGEN 79254), and three replicates were arranged. The RNA-seq libraries were prepared using the NEBNext PolyA mRNA Magnetic Isolation Module (NEB E7490S), NEBNext RNA First Strand Synthesis Module (NEB E7525S), NEBNext mRNA Second Strand Synthesis Module (NEB E6111S) and NEBNext Ultra DNA Library Prep Kit for Illumina (NEB E7370L). All samples were further subjected to NGS analysis using the Illumina HiSeq 4000 platform.

#### Bulk RNA-seq data processing

In the bulk RNA-seq library, the sequencing reads were aligned to the human reference genome (GRCh37/hg19) using HISAT2 (2.0.4)^33–35^ and assembled and quantified by StringTie (1.3.5)^33,36^.

### Single-cell RNA-seq and data analysis

#### Single-cell library preparation

The K562 cells were infected with the validated pgRNA hotspot_10_25-pg2. Eight days post lentiviral infection, the pgRNA-infected cells were FACS-sorted for single-cell library preparation. The single-cell library was prepared using the established protocol described previously^23^. Briefly, polyA+ RNA was reverse transcribed through tailed oligo-dT priming directly in whole cell lysate (single droplet) using Moloney Murine Leukemia Virus Reverse Transcriptase (MMLV RT) and temperature switch oligos. The resulting full-length cDNA contains the complete 5′ end of the mRNA, as well as an anchor sequence that serves as a universal priming site for second strand synthesis. The cDNA was pre-amplified using 15 cycles with Kapa HiFi Hotstart Readymix, then tagmented at 55°C for 5 min in 20 μl reaction following Illumina Nextera DNA preparation kit. 35 μl of PB was added to the tagmentation reaction mix to quench the reaction, and the tagmented DNA was purified with 20 μl of Ampure beads (sample to beads ratio of 1:0.6). Purified DNA was then amplified by 12 cycles of standard Nextera PCR. The prepared libraries were sequenced on Illumina HiSeq4000.

#### Single cell RNA-seq processing

The single cell RNAseq data were processed using the Drop-seq pipeline developed by the McCarroll lab^23^. The low quality reads (lower than Q10) and PCR duplicates were removed. Cells were decreasingly ranked by the total number of read counts. Cells ranked before the inflection point of the cumulative distribution were selected for the following analysis. Each cell was first normalized by counts per million (CPM). The value *E*_*i*_ was computed as the sum of CPMs for a given gene across all the cells. The *E*_*total*_ was calculated as the sum of *E*_*i*_ for all the genes. Then a *P*_*i*_ was computed as *P*_*i*_ =*E*_*i*_ / *E*_*total*_. In a given cell *j*, the normalized gene expressions of all genes were assumed to follow a binomial distribution independently identically *G*_*i j*_*∼B* (*N*_*j*_, *P*_*i*_), where *G*_*i j*_ is the expected read of gene *i* in cell *j, N*_*j*_ is the total read for cell *j*. A p-value was computed to evaluate how each gene expression in each cell significantly deviated from the expectation based on the binomial distribution. We also calculated p-values for genes in the negative control (Δ*AAVS1*) and wild type bulk RNA-seq data in the same way.

### Single-cell trajectory branching and pseudotime analysis

Because hotspot deletion severely deteriorated cell proliferation, we focused on analyzing the apoptosis genes annotated in the KEGG database ^37^. The 99 apoptosis genes that showed differential expression upon deleting hotspot_10_25 (*chr10: 74,123,469-74,124,868*) in at least 10% ∼ 15% cells (*p*-value < 0.05) were selected. All the normalized single cells and bulk data were clustered with trajectory branching and pseudotime analysis using Monocle^24,38^. Monocle assigned a specific pseudotime value and a “State” to each cell. Cells with the same “State” and similar pseudotime were clustered together^38^ and then the relative gene expression in each cluster was computed.

### Differentially expressed genes identified from pseudotime analysis

To identify differentially expressed genes (DEGs) pairwisely between different stages, in the pseudotime analysis, a *Wilcoxon* Rank-Sum Test^39^ was used to identify genes significantly up- and down-regulated in the cell states pair.

### Validation of synthetic lethal pairs by cell proliferation assay

#### Selection of the targeting sgRNA for each gene

To explore the synthetic lethal pairs among the four significant down-regulated genes located within the same TAD of Hotspot_10_25 after the hotspot deletion, we first determined the targeting sgRNA to ensure efficient knock down of each gene. Three sgRNAs were selected to target the promoter region of each gene from the hCRISPRi-v2 library^19^, and a non-targeting sgRNA was set as a control. These sgRNAs were further cloned into the lentiviral expression vector with an EGFP selection marker, then were transduced into K562 cells stably expressing KRAB-dCas9 protein through lentiviral infection. Three days post infection, the EGFP positive cells were FACS-sorted, and the total RNA of each sample was extracted using RNeasy Mini Kit (QIAGEN 79254). The cDNAs were synthesized from 2 μg of total RNA using Quantscript RT Kit (TIANGEN KR103-04), and real-time qPCR was performed with TB Green™ Premix Ex Taq™ II (Tli RNaseH Plus, TAKARA) to detect the expressions of each indicated gene as well as the reference gene *GAPDH*. The sgRNAs showing the most significant knock-down effect were selected for subsequent experiments to evaluate the synergistic effect. All the primers used in real-time qPCR were listed in **Supplementary Table 5**.

### Evaluation of the growth effect of each individual gene and gene pair in K562 cells

The above selected four sgRNAs were grouped into six gRNA-pairs targeting six gene-pairs. The four sgRNAs and six pgRNAs were respectively cloned into the lentiviral expression vector with an EGFP selection marker, then were transduced into K562 cells stably expressing KRAB-dCas9 protein at an MOI of < 1. The cell proliferation assay was performed as described above. The first time point of FACS analysis was at 6 days post lentiviral infection, and the experiment lasted for another 12 days.

## Notes

### Competing Interest Statement

The authors have declared no competing interest.

